# Genomic re-assessment of the transposable element landscape of the potato genome

**DOI:** 10.1101/701888

**Authors:** Diego Zavallo, Juan Manuel Crescente, Magdalena Gantuz, Melisa Leone, Leonardo Sebastian Vanzetti, Ricardo Williams Masuelli, Sebastian Asurmendi

## Abstract

Transposable elements (TEs) are DNA sequences with the ability to auto-replicate and move throughout the host genome. TEs are major drivers in stress response and genome evolution. Given their significance, the development of clear and efficient TE annotation pipelines has become essential for many species. The latest *de novo* TE discovery tools, along with available TEs from Repbase and sRNA-seq data, allowed us to perform a reliable potato TEs detection, classification and annotation through an open-source and freely available pipeline (https://github.com/DiegoZavallo/TE_Discovery). Using a variety of tools, approaches and rules, our pipeline revealed that ca. 16% of the potato genome can be clearly annotated as TEs. Additionally, we described the distribution of the different types of TEs across the genome, where LTRs and MITEs present a clear clustering pattern in pericentromeric and subtelomeric/telomeric regions respectively. Finally, we analyzed the insertion age and distribution of LTR retrotransposon families which display a distinct pattern between the two major superfamilies. While older Gypsy elements concentrated around heterochromatic regions, younger Copia elements located predominantly on euchromatic regions. Overall, we delivered not only a reliable, ready-to-use potato TE annotation files, but also all the necessary steps to perform *de novo* detection for other species.

**Key Message:** We provide a comprehensive and reliable potato TE landscape, based on a wide variety of identification tools and integrative approaches, producing clear and ready-to-use outputs for the scientific community.

## Introduction

Transposable elements (TEs) are DNA sequences with the ability to auto-replicate and move throughout the genome. In plants, TEs can occupy a large proportion of the genome, representing more than half of the total genomic DNA in some cases. For example, they comprise about 85% and 88% of wheat and maize genomes, respectively (Schnable et al., 2009; Appels et al., 2018) which may indicates the relevance of these elements to genome architecture and size (Roessler et al., 2018).

Furthermore, over the past few years an increasing number of studies have shed light on their importance in gene regulation (Hirsch and Springer, 2017; Judd and Feschotte, 2018) stress response and genome evolution (Hosaka and Kakutani, 2018).

TEs are usually divided into two major classes based on their mechanism of transposition. Class I elements (or retrotransposons) propagate via a reverse-transcribed RNA intermediate, whereas class II elements (or DNA transposons) move through a “cut and paste” mechanism. Another type of classification is by their ability to transpose on their own (i.e. autonomous), characteristic shared by some TEs from both classes. The non-autonomous elements can move but rely on autonomous TEs for their mobility (Wicker et al., 2007).

Given their significance, the identification, classification and annotation of TEs has emerged as a new field of great interest in science, which involves both wet-lab biology and bioinformatics. As Hoen et al. (2015) describe in their review on TE annotation benchmarking, precise detection and annotation of TEs is a difficult task due to their great diversity, both within and among genomes. TEs differ across multiple attributes, including transposition mechanism, sequence, length, repetitiveness, and chromosomal distribution. In addition, whereas recently inserted TEs have a relatively low variability within the family, over time they accumulate mutations and diverge, making them harder to detect (Hoen et al., 2015).

There are two main strategies for TE annotation: homology-based and *de novo* identification, which can also be referred to as library-based and signature-based, respectively. The homology-based strategy uses libraries of known TEs such as the Repbase repository (Jurka et al., 2005) to screen genomes in order to identify similar sequences, most commonly by using RepeatMasker (Smit et al., 1996). On the other hand, *de novo* approaches use characteristic structural features, such as LTRs (Long Terminal Repeats) for retrotransposons and TIRs (Terminal Inverted Repeats) for DNA transposons, to identify new elements. Moreover, autonomous TEs have conserved structures like RT (reverse transcriptase) or TR (transposase) that can also be used for accurate TE identification (Wicker et al., 2007). Several tools based on structural features, such as LTRharvest (Ellinghaus et al., 2008) and TIRvish command, which are part of the GenomeTools suite (Gremme et al., 2013), are available. Other tools based on this criteria are specifically designed to discover different types of TE families (e.g. SINE Scan (Mao and Wang, 2016), HelitronScanner (Xiong et al., 2014) and MITE Tracker (Crescente et al., 2018)). Another *de novo* based strategy relies on the most important biological mechanism that silences TEs - RNA directed DNA Methylation (RdDM) - in which double-stranded RNAs (dsRNAs) are processed into 21-24 nt small interfering RNAs (siRNAs) and guide methylation on homologous DNA loci. For instance, TASR (for Transposon Annotation using Small RNAs) tool uses sRNAs Illumina data as guide for TE annotation/identification (El Baidouri et al., 2015). However, complete and accurate TEs annotation will likely require a combination of both homology-based and *de novo* methods together with an additional manual curation step (Platt et al., 2016).

Nevertheless, all of these tools often give a representative sequence for each family (usually a full-length TE sequence), but fail presenting their copies across the genome, not only for other potentially autonomous copies, but also for members that are partially or entirely deficient in one or more domains. Furthermore, in the case of non-autonomous TEs (e.g. SINEs, TRIMs and MITEs), the amount of copies across the genome is high due to their repetitive content and their short length; however this information is usually not presented.

For the scientific community, especially in genomics, the need for reliable annotation is becoming fundamental. Generally, for each genome sequencing project, annotations consisting mainly of protein-coding genes (structural and functional) and miRNAs genes are made, whereas TEs remain poorly annotated As an example, the Gramene Database (http://www.gramene.org) is a curated, open-source database for comparative functional genomics in crops and model plant species with information on more than two million genes from 67 plant genomes. By contrast, the PGSB Repeat Element Database (Nussbaumer et al., 2012) which compiles publicly available repeat sequences from TREP (Wicker et al., 2002), TIGR repeats (Ouyang and Buell, 2004), PlantSat (Macas et al., 2002), Genbank and *de novo* detected LTR retrotransposon sequences from LTR_STRUC (McCarthy and McDonald, 2003) only comprises 62.000 sequences. Nonetheless, in the latest version of sunflower genome, Badouin et al. (2017) performed a comprehensive search for repeat elements by developing a new tool called Tephra (https://github.com/sestaton/tephra) (Staton, 2018), which discovers and annotates all types of transposons. Tephra combines existing specific transposon discovery tools for all types of TEs, classifies and annotates them, but still lacks information on copy numbers across the genome. A recent study addressed this issue by applying a method called “Russian doll” due to its nesting strategy. This method builds nested libraries establishing different search rules for each one of them. The first one includes only “potentially autonomous TEs”, the second one contains “total TEs”, including non-autonomous and a third one that also includes uncategorized “repeated elements” (Berthelier et al., 2018).

*Solanum tuberosum*, the cultivated potato, is the third most important food crop after rice and wheat, and the main horticultural crop (Devaux et al., 2014). The sequencing of the *S. tuberosum* genome resulted in an assembly of 727 Mb of 810.6 Mb sequenced. Because most potato cultivars are autotetraploid (2n=4x=48) and highly heterozygous, sequencing was performed on a homozygous doubled-monoploid potato clone. The latest potato genome assembly (4.03) contains 39.031 annotated genes and 62.2% of the genome corresponds to repetitive elements at scaffold level (Consortium et al., 2011).

Many attempts have been made to discover repetitive elements in Solanaceae families. Most of these studies were mainly focused on tandem repetitive elements, whereas studies of complex repetitive elements were mostly performed on limited groups of TEs (Mehra et al., 2015). The researchers assessed the complex repetitive elements in potato *(S. tuberosum*) and tomato *(S. lycopersicum)* genomes, identifying 629,713 and 589,561 repetitive elements, respectively. Mehra et al. used RepeatModeler (http://repeatmasker.org/RepeatModeler.html), which employs a repetitiveness-based strategy, and enriched the amount of repeat families previously identified in the 4.03 version of the potato genome with RepeatMasker (Smit et al., 1996).

In this study we present an optimized pipeline of transposable elements detection and annotation from *S. tuberosum*. Our strategy relies on the combination of the latest *de novo* TE identification tools, available TEs from Repbase and Illumina sRNA-seq data to obtain TEs. We then find copies and applied a series of filters depending on the TE family to obtain a comprehensive and curated whole-genome atlas of potato transposable elements. Furthermore, we provide to the research community our pipeline, annotation results and files, which are publicly available at https://github.com/DiegoZavallo/TE_Discovery to encourage reproducibility and the eventual implementation of our framework in diverse organisms.

## Materials and Methods

### Input data

The latest assembly version of *Solanum tuberosum* genome sequence was downloaded from the Potato Genome Sequencing Consortium (PGSC v4.03 of the doubled monoploid *S. tuberosum* Group Phureja DM1-3). Potato TEs sequences from Repbase Giri (Jurka et al., 2005) were downloaded prior registration. Note that LTRs transposons are divided in LTR (Long Terminal Repeat) and I (Internal) sequences, hence must be concatenated. We gathered 126 LTRs, 18 LINEs, 2 SINEs and 42 TIRs family sequences (https://www.girinst.org/repbase/update/search.php?query=tuberosum\&querytype=Taxonoy).

Illumina sRNA-seq data for TASR run was generated by our lab (data unpublished); however, any available data from public repositories such as SRA (https://www.ncbi.nlm.nih.gov/sra) could be used as input.

### Transposable elements identification

TEs obtained from different sources were merged together according to each element classification. Tephra, which uses several structure-based tools, was applied to harvest different kind of TEs in the potato genome. *Tephra all* command (which runs all subcommands) was executed using tephra config.yml file with default configuration parameters with the exception of *S. tuberosum* TEs from Repbase for the repeatdb parameter. A total of 1,325 Helitrons, 7,694 LTRs, 2,994 MITEs, 2,414 TIRs and 7,011 TRIMs families were found with this program. No non-LTR TEs (LINEs and SINEs) were found by Tephra. TASR (El Baidouri et al., 2015), a tool for de novo discovery of TEs using small RNA data, was also used in this work. 21, 22 and 24 nt sRNAs were parsed from files belonging to all treatments and replicates from our sRNA-Seq data and subsequently concatenated to be used as input. TASR.v.1.1.pl perl script was run with default parameters except for: *-cpu 14, -nsirna 10, and -cnumber 5*. A total of 1,916 families were found and presented as multifasta files comprising all the elements for each family. A perl script provided by TASR developer was run in order to generate a consensus sequence for each family. Then we created a single multifasta file with the entire consensus. The PASTEC tool (Hoede et al., 2014) was used for this purpose, since TASR does not classify TEs into categories. From the 1,916 families discovered, a total of 891 LTRs, 49 LINEs, 15 SINEs, 9 TRIMs, 2 LARDs, 84 TIRs, 35 MITEs and 5 Helitrons were classified. For MITEs discovery, MITE Tracker (Crescente et al., 2018) was employed using default parameters. A total of 1,045 MITEs elements were detected. SINE_Scan (Mao and Wang, 2016), an efficient de novo tool to discover SINEs was also used in this work with default parameters. A total of 13 SINEs families were detected. As a result, we obtained 8,711 LTRs elements from Tephra, TASR and Repbase, 67 LINEs elements from TASR and Repbase, 30 SINEs elements from TASR, Repbase and SINE_Scan, 540 TIRs elements from Tephra, TASR and Repbase, 4,074 MITEs elements from Tephra, TASR and MITE Tracker and 1,330 Helitrons elements from Tephra and TASR.

### Pipeline description

To detect, filter and annotate TEs copies across the potato genome the obtained TEs list was subjected to an in house pipeline. The pipeline was developed using bash scripts and Jupyter notebooks.

*1*.*1 add annotation*.*ipynb* uses TEs sequences retrieved from each program and merge them together into one multi-fasta file per studied TE type (LTRs, LINEs, SINEs, TRIMs, LARDs, TIRs, MITEs and Helitrons). This script adds to each sequence a unique identifier containing an auto-incremented number, TE classification and the program source from which it was obtained.

*2*.*1 vsearch*.*sh* uses VSearch program to cluster similar sequences that share 80% identity, according to the 80/80/80 rule in the study of Wicker et al. (2007). It is executed once per TE type, thus obtaining as a result TEs clustered by type.

*2*.*2 vsearch merge*.*ipynb* script uses vsearch outputs to create fasta files containing one TE per family. It also adds a family description indicating which program the members came from.

*3*.*1 blast*.*sh* performs a genome-wide BLASTn search using the files from the previous step and searches for TEs in the potato genome. For this task, the script uses the following parameters: *-perc identity 80, -evalue 10e-3 and -task blastn* (except for LTRs). *-qcov hsp perc 80* was used for SINEs, TRIMs and MITEs, whereas *-qcov hsp perc 50* was set for the rest.

*3*.*2 blast filter*.*ipynb* filters BLASTn results by using parameters according to Table 1. First, each file is filtered by a length range defined by min len and max len. Afterwards, a length threshold range is calculated by multiplying the query length by a min and max subject length subject length. The subject sequence has to be inside this range to be considered valid. Later, minimum identity percentage and query coverage are required for the sequences to be valid. Finally, duplicated hits are removed by searching those whose start and end positions overlaps within a margin of plus or minus 5 nt.

**Table 1:** TEs filtering parameters. Parameters established to filter copies element for “Autonomous TEs” and “All TEs” which includes autonomous and non-autonomous elements. *min-len*: minimum element length; *max-len*: maximum element length; *min-pid*: minimum query identity percentage; *min-threshold*: minimum threshold length between query and subject; *max-threshold*: maximum threshold length between query and subject; *min-cov*: minimum query coverage percentage; *overlap*: margin of plus or minus nucleotides overlap between query and subject to be considered as duplicates.

| Family | min-len<br>(nt) | max-len<br>(nt) | min-pid<br>(%) | min-threshold<br>d (ratio) | max-threshold<br>d (ratio) | min-cov (%) | overlap<br>(nt) |
| --- | --- | --- | --- | --- | --- | --- | --- |
| **Autonomous TEs** |  |  |  |  |  |  |  |
| LTR | 650 | ∞ | 95 | 0.9 | 1.1 | 95 | 5 |
| LINE | 1500 | ∞ | 95 | 0.9 | 1.1 | 95 | 5 |
| TIR | 700 | ∞ | 95 | 0.9 | 1.1 | 95 | 5 |
| Helitron | 2000 | ∞ | 95 | 0.9 | 1.1 | 95 | 5 |
| **All TEs** |  |  |  |  |  |  |  |
| LTR | 650 | ∞ | 80 | 0.5 | 1.5 | 50 | 5 |
| LINE | 1500 | ∞ | 80 | 0.5 | 1.5 | 50 | 5 |
| SINE | 150 | 800 | 80 | 0.8 | 1.2 | 80 | 5 |
| TRIM | 600 | ∞ | 90 | 0.9 | 1.1 | 90 | 5 |
| LARD | 4000 | ∞ | 80 | 0.5 | 1.5 | 50 | 5 |
| TIR | 700 | ∞ | 80 | 0.5 | 1.5 | 50 | 5 |
| MITE | 50 | 800 | 80 | 0.85 | 1.15 | 90 | 5 |
| Helitron | 2000 | ∞ | 80 | 0.5 | 1.5 | 50 | 5 |

*4*.*1 annotate*.*ipynb* transforms the BLAST tab-delimited results to a *gff3* format file, adding a detailed description for each TE. The description includes TE id (a numeric identifier of the element after clustering) source name (original id name of the element before clustering) type (family type of the element), source (program or tool from which the TEs were detected) and unique id (unique identifier for copy element).

### LTR age

To determine tentative LTRs insertion age we used *tephra ltrage* command implemented by the Tephra package. This command uses the Tephra-discovered full length LTRs, aligns the LTR sequences and generates a neighbor-joining guide tree with MUSCLE (Edgar 2004). The alignment and guide tree are used to generate an alignment in PHYLIP format. A likelihood divergence estimate was calculated with baseml from PAML (Yang 2007) by specifying the K80 substitution model. This divergence value (hereafter d) was used to calculate LTR-RT age with the formula T = d/2r, where r= 1e8 is the default substitution rate.

### Data resource

Scripts from this work, including all pipeline steps as well as circos ideogram, distance histogram and LTR age plot scripts are available at (https://github.com/DiegoZavallo/TEDiscovery). Annotation and fasta files are available as supporting information.

## Results and Discussion

The goal of the present work was to assemble a comprehensive TE repertoire of the potato genome and provide legible, ready-to-use files for the scientific community. To address this issue, we used a combination of different approaches to identify TEs such as similarity-based, structure-based and mapping-based strategies. Moreover, we introduce a set of scripts to gather, detect, filter and ultimately annotate TE copies from all classes across the potato genome and present *gff3* files of TE features. Additionally, we display the accumulation and distribution across the genome of the different types of TEs and a table summarizing data and metrics, such as distances to nearest gene and LTRs insertion ages.

### TEs in the potato genome

Public *S. tuberosum* Repbase library (Jurka et al., 2005), the potato reference genome and Illumina small RNA-seq data were used as **input data. TEs family detection** was executed by applying two “All TEs” tools: Tephra and TASR, which discover *de novo* TEs with structure-based and map-based approaches, respectively. To complement the search, we applied “Specific TEs” tools: MITE Tracker and SINE_Scan. The obtained sequences were merged into multi-fasta files, one for each type of TE, and headers were renamed. A **clustering** step was carried out with Vsearch to reduce redundancy (Online Resource 1). Next, **detection of copies** in the genome of the different TE families was achieved by conducting a BLAST search with specific parameters according to the type of TE (see Materials and Methods section). A **copy filter** step was implemented by establishing several rules with very astringent criteria to detect “potential autonomous TEs” and a second set of rules for “All TEs” with more relaxed parameters to account for autonomous and non-autonomous TEs of all types (Table 1). Finally, an **annotation** step was performed to generate *gff3* files containing a description for each element that includes TE type and the detection tool that identified it. Figure 1 shows an overview of the pipeline used to detect and re-annotate the potato TEs.

**Figure 1.**
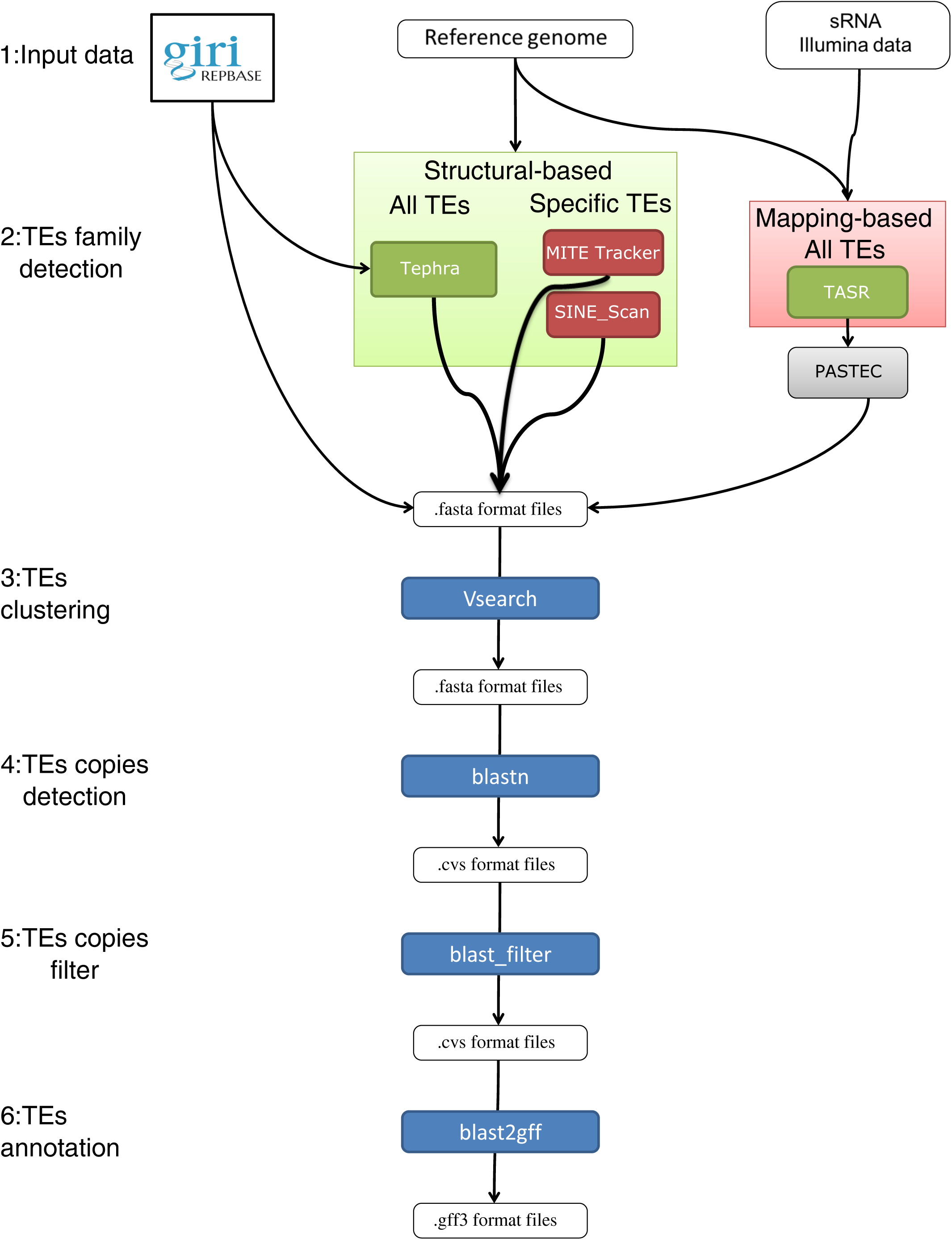
Overview of the TE Discovery pipeline. *1_Input data*: last genome assembly, available TE sequences from Repbase Giri and sRNA-seq Illumina data are used as input data. *2_TEs family detection*: the detection of putative TEs is performed by using four tools combining two detection approaches. The resulted sequences are merged into multi-fasta files for each type of TEs. *3_TEs clustering*: Each TE type sequences are clustered with Vsearch to reduce redundancy. *4_TEs copy detection*: blastn is performed to the clustered sequences with specific parameters according to each type of TE to detect copies across the genome. *5_TEs copy filter*: the detected copies are subjected to filtering steps to detect “potential autonomous TEs” and/or “All TEs” including non- autonomous TEs. *6_TEs annotation*: TEs annotation *gff3* files are generated for each type of TEs with detailed descriptions and merged into one single file.

TE content is highly variable in plants and usually displays a positive correlation with genome size. For instance, as much as 85 % of maize genome or 70 % of Norway spruce genome (Nystedt et al., 2013) has been annotated as transposons including unclassified ones, whereas in the more compact *Arabidopsis thaliana* genome TE content is only 21 % (Ahmed et al., 2011). In potato, the data presented by Mehra et al. (2015) comprised an annotation file of 1,061,377 repetitive elements, including rRNA, tRNA, simple repeats and low complexity elements which represents almost 50% of the genome. When only the most complex elements (i.e. transposons) were taking into account, the coverage percentage dropped to nearly 34%.

However, our pipeline revealed a TE content of ∼16% (excluding the unanchored ChrUn), representing half of the genomic coverage according to the data presented by Mehra et al. (2015).

Of those ∼16%, LTRs comprised around 13% of the potato genome, which corresponds to over 80% of the total TEs. The most abundant superfamily was Gypsy, whereas the other types of TEs barely made up 1% of the genome coverage. For instance, each DNA TE (TIRs, MITEs and Helitrons) covered almost the same percentage of the genome with 0.51, 0.72 and 0.72 %, respectively. These coverage ratio patterns are in agreement with results from most plant genomes that have TE identification projects (Du et al., 2010; Andorf et al., 2016; Badouin et al., 2017; Alaux et al., 2018). Table 2 summarizes the amount and diversity of all identified TEs, filtered copies and proportion in the genome of all TE families.

**Table 2:** Quantity and diversity of all identified TEs. Number of TE families, number of TE copies and TEs genome coverage (%) of all types of TEs.

| Class | Order/Family | TEs identified | TEs copies | Genome coverage (%) |
| --- | --- | --- | --- | --- |
| I | LTR/Copia | 2,541 | 13,044 | 2.46 |
| I | LTR/Gypsy | 5,736 | 170,380 | 10.68 |
| I | LTR/Unclassified | 29 | 604 | 0.06 |
|  | Total LTRs | 8,306 | 184,028 | 13.2 |
| I | LINE | 65 | 248 | 0.1 |
| I | SINE | 24 | 2,477 | 0.07 |
| I | TRIM | 6,908 | 13,491 | 0.67 |
| I | LARD | 2 | 18 | 0.002 |
| **Total Class I** |  | 15,305 | 200,262 | 14.04 |
| II | TIR/hAT | 112 | 382 | 0.05 |
| II | TIR/Mariner | 1,466 | 1,591 | 0.29 |
| II | TIR/Harbiner | 16 | 219 | 0.02 |
| II | TIR/Mutator | 575 | 941 | 0.13 |
| II | TIR/CACTA | 131 | 117 | 0.02 |
| II | TIR/Unclassified | 3 | 110 | 0.01 |
|  | Total TIRs | 2,303 | 3,380 | 0.51 |
| II | MITEs | 3,515 | 38,205 | 0.72 |
| II | Helitrons | 1,322 | 1,163 | 0.72 |
| **Total Class II** |  | 7,140 | 42,748 | 1.99 |
| **Total TEs** |  | 22,445 | 243,010 | 16.03 |

To understand more deeply the discrepancy between the coverage of the TE genome presented here and the one reported by Mehra et al. (2015), we should observe other differences between both works. For instance, even though we describe all types of TE families reported by Wicker et al. (2007), including LARDs, TRIMs and MITEs, which were absent in Mehra study, we annotated less than half of the sequences. We applied a clustering method to decrease redundancy and established a set of filters selected specifically for each type of TEs, which, is aimed to reduce false signal. Moreover, as already mentioned, Mehra et al. used a unique tool that relies on repetitiveness-based strategy, leaving aside a wide variety of detection methods that in this work we have combined, which is indicated to improve TE detection efficiency (Kamoun et al., 2013; Hoen et al., 2015; Arensburger et al., 2016).

We scored a total of 243,010 elements compared to 629,713 complex elements previously found by Mehra et. al. and when compared, 198,025 (81%) of our sequences overlapped with their set, which evidences the effectiveness of both annotation pipelines.

To test the effectiveness of the pipeline, we run it on a well annotated genome such as soybean. SoyTEdb represents an example of a thoroughly annotated TE database in which a combination of structure-based and homology-based approaches was used to structurally annotate and clearly categorize TEs in the soybean genome (Du et al., 2010). The authors reported over 38,000 TEs representing ∼17% of the genome. However, when they informed the genome coverage they included fragments defined by RepeatMasker (i.e. low complex repeats) rising up to 58% of the soybean genome. These data may indicate the existence of a large set of repetitive sequences in the genome that cannot be annotated as TE with current knowledge about the structure of TE. One hypothesis that could arise from this is that the structural patterns that represent these non-annotated TEs are not yet thoroughly described, or they are just repetitive DNA that simply cannot be assigned as TEs. We used the 38,000 annotated TEs to run our pipeline for copy elements discovery and filtering steps and we came out with similar genomic coverage (∼23%) and more than 75% of the TEs sequences overlapped. Even though this was only a test, since the full run of our pipeline on another species would require a more comprehensive approach which exceeds this work, it validates that this pipeline does not underestimate the occurrence of TEs in the genome.

Finally, we deliver ready-to-use annotation files in *gff3* format of all TEs annotated with our pipeline with detailed descriptions in the ninth column including *TE id, source name, type, source* and *unique id* (Online Resource 2).

### Distribution of TEs across the potato genome

It is well know that the distribution, amount and genome coverage of TEs vary greatly, particularly between plants and animals, where LTRs and non-LTRs (LINEs and SINEs) are the predominant type of TE, respectively (Chalopin et al. 2015). Moreover, TE distribution is highly dependent on the family type. Some TEs are more prone to concentrate in regions near protein coding genes while others are more equally distributed along the genome.

To assess the type-specific landscape of the diverse TE categories, we performed circos ideograms for each TE type separated by class, as well as gene density as reference. Each concentric circle represents the coverage percentage of one type of TE. Since they have different coverage ranges, each type has its own color pallet to appreciate the distribution across the chromosomes (Figure 2).

**Figure 2.**
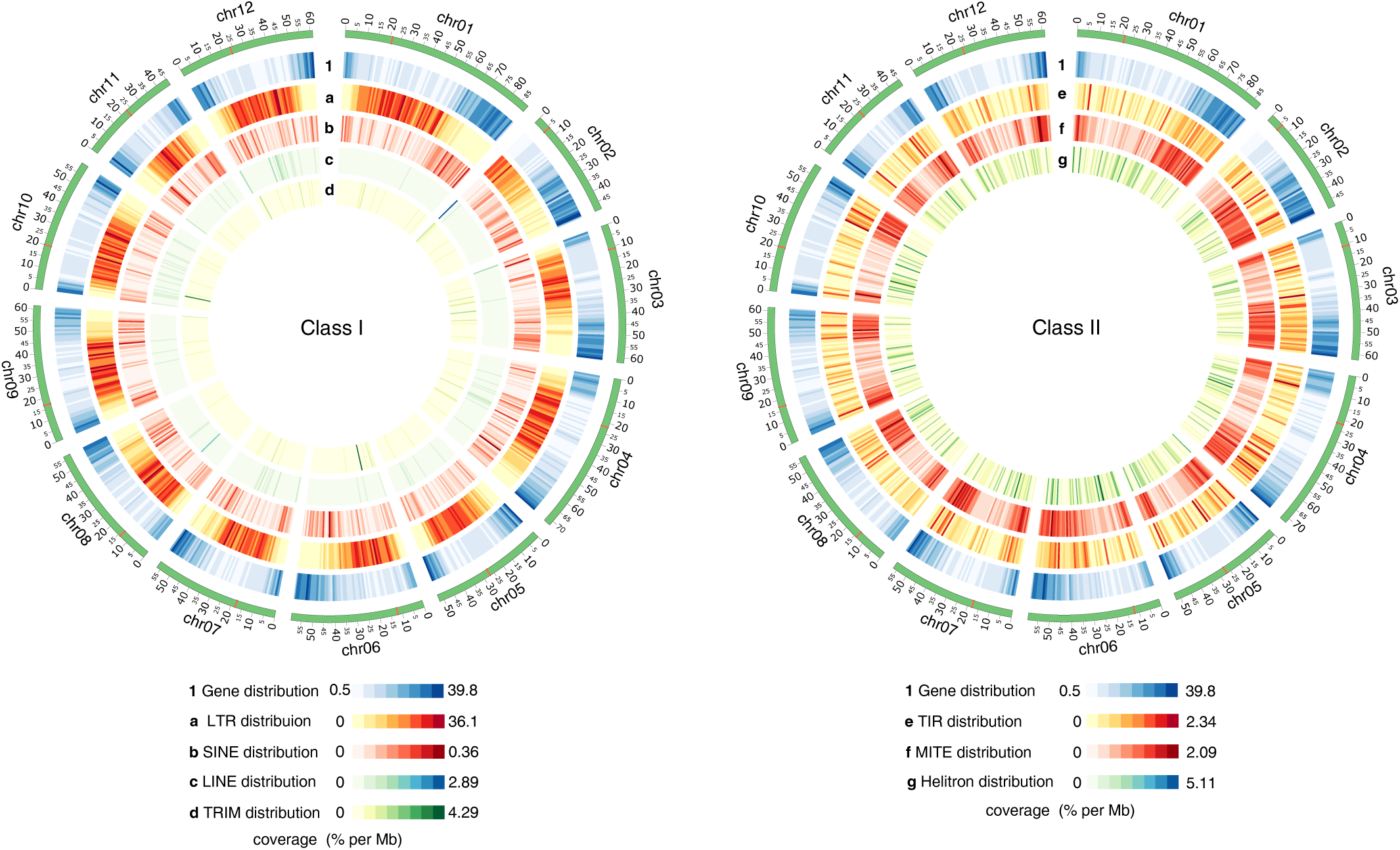
Comprehensive circos ideograms of TEs of the potato genome. Left panel: Retrotransposon TEs (Class I). Right panel: DNA TEs (Class II). Each concentric circle represents a different type of TE with their own color pallet and range of coverage so the distribution across the chromosomes can be appreciated. Each line within the circles represents coverage percentage per Mb. (1) Gene distribution. (a) LTR distribution. (b) SINE distribution. (c) LINE distribution. (d) TRIM distribution. (e) TIR distribution. (f) MITE distribution. (g) Helitron distribution.

Left panel of Figure 2 shows the class I elements, where LTRs stand out, not only for their clear pattern of clustering around centromeric and pericentromeric regions, but also for their high coverage in some areas of the chromosomes. For instance, each dark red line of the second circle represent up to 36% of LTRs coverage per Mb, which explains the 13% genome-wide coverage for this kind of TE (Table 2). Furthermore, LTRs distribution is virtually opposite to protein coding gene distribution (Figure 2, left panel). This behavior has already been reported for other plants (Baucom et al., 2009; Paterson et al., 2009; Badouin et al., 2017).

Conversely, SINEs, which are the least represented type (besides LARDs which are not shown since we discovered only 18 elements) seem to have an even distribution pattern with a slight tendency towards telomeric and subtelomeric regions in some chromosomes. LINEs and TRIMs also display homogeneous distribution patterns with some coverage hotspots, mainly near telomeric regions. Due to their length and filtering parameters established, only 248 LINEs were found in the genome using our methodology, contrasting with more than 50,000 elements found by Mehra et al. (2015). Given that Heitkam et al. (2014) extracted 59,390 intact LINE sequences from 23 plant genomes in order to classify them into families, it is somehow unlikely that potato alone could have 50,000 LINEs. A look at the data presented by Mehra et al. (2015) shows that some elements annotated as LINEs are small fragments with some identity to LINEs given by the RepeatMasker tool which in our case were removed by the filtering process.

In a recent work, Gao et al. (2016) performed a comprehensive analysis of TRIMs in 48 plant genomes, including *S. tuberosum*. They observed that TRIMs are generally enriched in genic regions and likely play a role in gene evolution (Gao et al., 2016). They discovered 12,473 copies in the potato genome representing 0.46% of the genome, which is consistent with our results (Table 2).

Right panel of Figure 2 shows class II TEs distribution in which MITEs display a clear concentration pattern around gene-rich subtelomeric regions. According to previous works, MITEs are often found close to or within genes, where they affect gene expression (Bureau and Wessler, 1994b). Indeed, MITEs may affect gene regulation via small RNA pathways, in addition to play the canonical role of TEs in the evolution by altering gene structure (Kuang et al., 2009; Gagliardi et al., 2019).

TIRs and Helitrons displayed an unbiased distribution across the chromosomes (Figure 2, right panel). Helitron chromosome distribution seems to vary by species. While in *Arabidopsis* Helitrons are enriched in gene-poor pericentromeric region, in maize they are more abundant in gene-rich regions (Yang and Bennetzen, 2009). Rice, on the other hand, exhibited a more erratic pattern of Helitron distribution (Yang and Bennetzen, 2009), more similar to our results.

In sum, this kind of analysis allowed us to have a holistic view of the different distribution patterns of TEs across the genome and to elucidate their potential role in transcriptional gene regulation.

### TEs and genes

The importance of TEs accumulation near genes, where these elements could influence gene expression, has been extensively reported (Bureau and Wessler 1994b,a; Wang et al., 2013). As we described above, we found a strong positive correlation between LTRs and pericentromeric regions as well as a positive correlation between MITEs and rich-gene subtelomeric regions. Hence, we determined the distances from the different types of elements to the closest gene in the genome in order to assess how many TEs are likely within or close to genes. For this purpose, we computed metrics such as median distance to the closest gene and percentage of TEs overlapping gene transcripts or near coding regions. LTRs were found to be generally far from genes with a median distance of 10.25 kb; 9.01% overlap with coding sequence genes and only 5.11% are in the immediate vicinity (up to 1 kb from the nearest gene), totaling a 32.23% within the first 5 kb, including elements within transcripts (Figure 3 and Online Resource 3). TRIMs exhibit a similar behavior, with a median distance of 8.12kb, 6.43% elements located within a gene and 6.19% in the range of 0-1 kb to the nearest transcript, comprising a 36.62% in the first 5 kb. In contrast, LINEs and SINEs are close to genes (median distance of 4.16 kb and 3.86 kb, respectively) with more than 20% elements within a transcript and rising up to >50% in the 0-5 Kb range (52.44% and 56.20% respectively).

**Figure 3.**
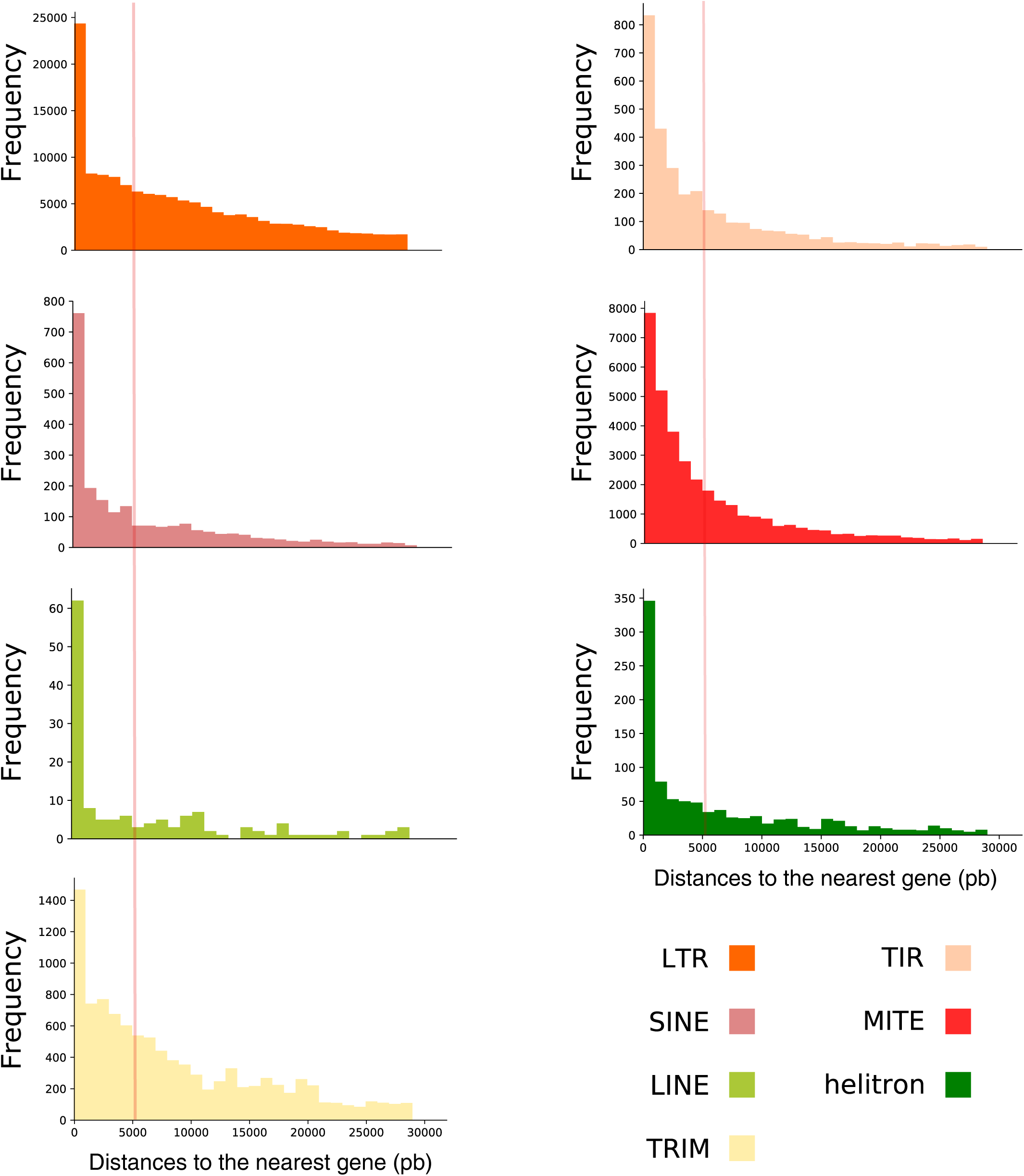
Frequency histograms of TE distance to the nearest gene. Left panel: Retrotransposon TEs (Class I). Right panel: DNA TEs (Class II). Bars on the left side of the red line represent the TE distance within the first 5kb to the nearest gene.

Class II TEs also have more than 50% of their elements within the first 5 kb. However, TIRs and MITEs distribution differs from that of Helitrons. TIRs and MITEs are the only types of TEs that appear to have less elements within a gene than in the 0-1 kb range (TIRs: 10.05% against 15.5% and a median distance of 3.32 kb; MITEs: 7.07% against 14.21% and a median distance of 3.53 kb). On the other hand, Helitrons display a similar pattern to LINEs and SINEs with a median of 4.56 kb and 23.65% elements within a transcript. These differences can be observed in the less pronounced curve of TIR and MITE histograms demonstrating that a significant proportion of elements are indeed in the proximity of genes but not necessarily within them. (Figure 3, right panel and Online Resource 3).

Overall, DNA TEs and nonLTRs retrotransposons have more than 50% of the elements inserted within transcripts or in a range of 0-5 kb from the nearest gene. Several studies have reported examples of transcriptional impact due to TEs insertion near genes in tomato (Xiao et al., 2008; Quadrana et al., 2014), potato (Momose et al., 2010; Kloosterman et al., 2013), melon (Martin et al., 2009) and orange (Butelli et al., 2012) among others. Several mechanisms including disruption of promoter or reduction of transcription through the spread of epigenetic silencing often suppress expression. However, TE can also introduce new sequences in the promoter, leading to up-regulation of proximal gene (Yan et al., 2004; Cowley and Oakey, 2013; Dubin et al., 2018). Insertion of a TE into the coding sequence can disrupt gene function, generally resulting in loss-of-function mutations, particularly if located in an exon. Intronic TEs can also be harmful, for instance by altering splicing patterns (Saze et al., 2013; Ong-Abdullah et al., 2015).

In contrast, LTRs and TRIMs located near genes barely exceed 30% and, with a few exceptions, they appear in intergenic, heterochromatic and gene-poor-regions (Kumar and Bennetzen, 1999). In *Arabidopsis*, Wang et al. (2013) reported that gene expression is positively correlated with the distance of the gene to the nearest TE, and negatively correlated with the number of proximal TEs. Whether LTR-like TEs specifically target these regions or if they are simply not selected against and accumulated in regions nearby genes remains unclear (Sigman and Slotkin, 2016).

### LTR elements age and evolution

LTRs are the most represented TEs in the majority of plant genomes, encompassing more than 75% of the nuclear genome of some species (Kumar and Bennetzen, 1999; Paz et al., 2017). For this reason, we decided to explore the evolutionary history of LTRs during potato genome evolution.

We analyzed the age and distribution of the elements discovered by Tephra (Staton, 2018) by means of *tephra ltrage* command that allowed the characterization of phylogenetic substructure within families of LTR retrotransposons. A total of 8,034 full-length LTRs were analyzed in terms of chromosomal distribution and insertion age by plotting all LTR elements together (Figure 4, upper panel), or grouped into superfamiles (Figure 4, bottom panel).

**Figure 4.**
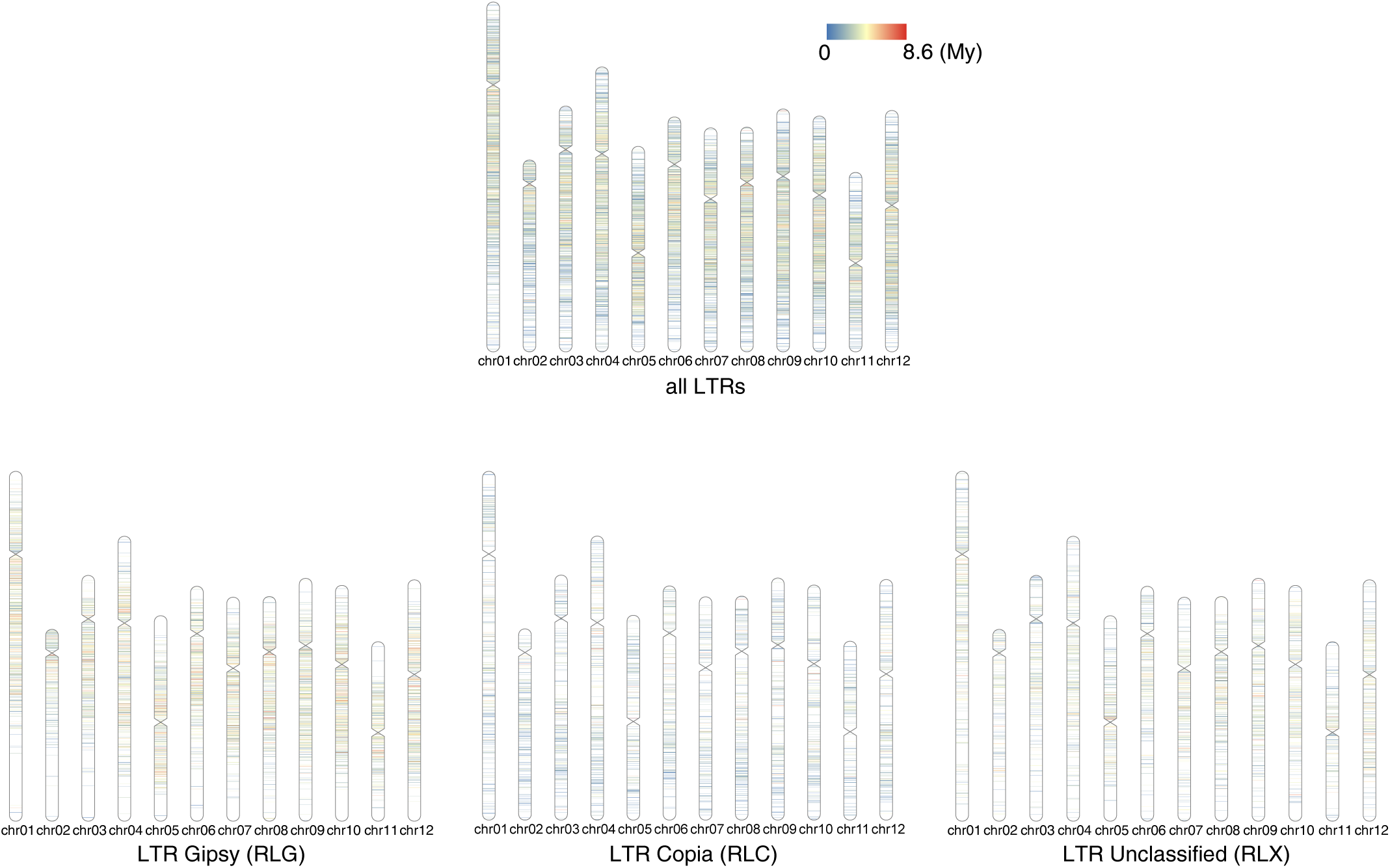
Chromosomal ideograms of LTRs age per family. Upper panel: chromosomal distribution of insertion age of all superfamilies of full-length LTR retrotransposons. Lower panel: chromosomal distribution of insertion age of Gypsy, Copia and Unclassified LTR retrotransposons separately.

Younger LTRs are enriched in euchromatic subtelomeric regions and correspond mainly to Copia (RLC) family of TEs (average insertion age of 2.54 mya), whereas older LTRs are more abundant in heterochromatic pericentromeric regions where Gypsy (RLG) elements are mostly located (average insertion age of 4.14 mya). These distributions are in agreement with previous findings in maize (Sun et al., 2018), wheat (Luo et al., 2017) and tomato (Paz et al., 2017).

Unclassified LTRs (RLX) display a more even distribution, although slightly towards to pericentromeric regions, as well as an intermediate insertion age (average insertion age of 3.78 mya) (Figure 4, bottom panel).

To determine whether these aging differences were a family trait or a genomic region-dependent mutation/substitution rate, we divided each chromosome in euchromatic and heterochromatic regions. In order to do so, we compared the pachytene karyotype previously published (Consortium et al., 2011) with the chromosomal ideograms we produced, and plotted the frequency of each LTR family by insertion age according to the karyotype determined region (Figure 5, upper panel). Gypsy family (RLG) display a Gaussian distribution centered around four millions years both in euchromatic and heterochromatic regions, whereas Copia family (RLC) have a chi-square distribution with a peak between two and three millions years on both regions. These results suggest that the insertion age of LTRs retrotransposons depends on the superfamily and not on differential mutation/substitution rate by region.

**Figure 5.**
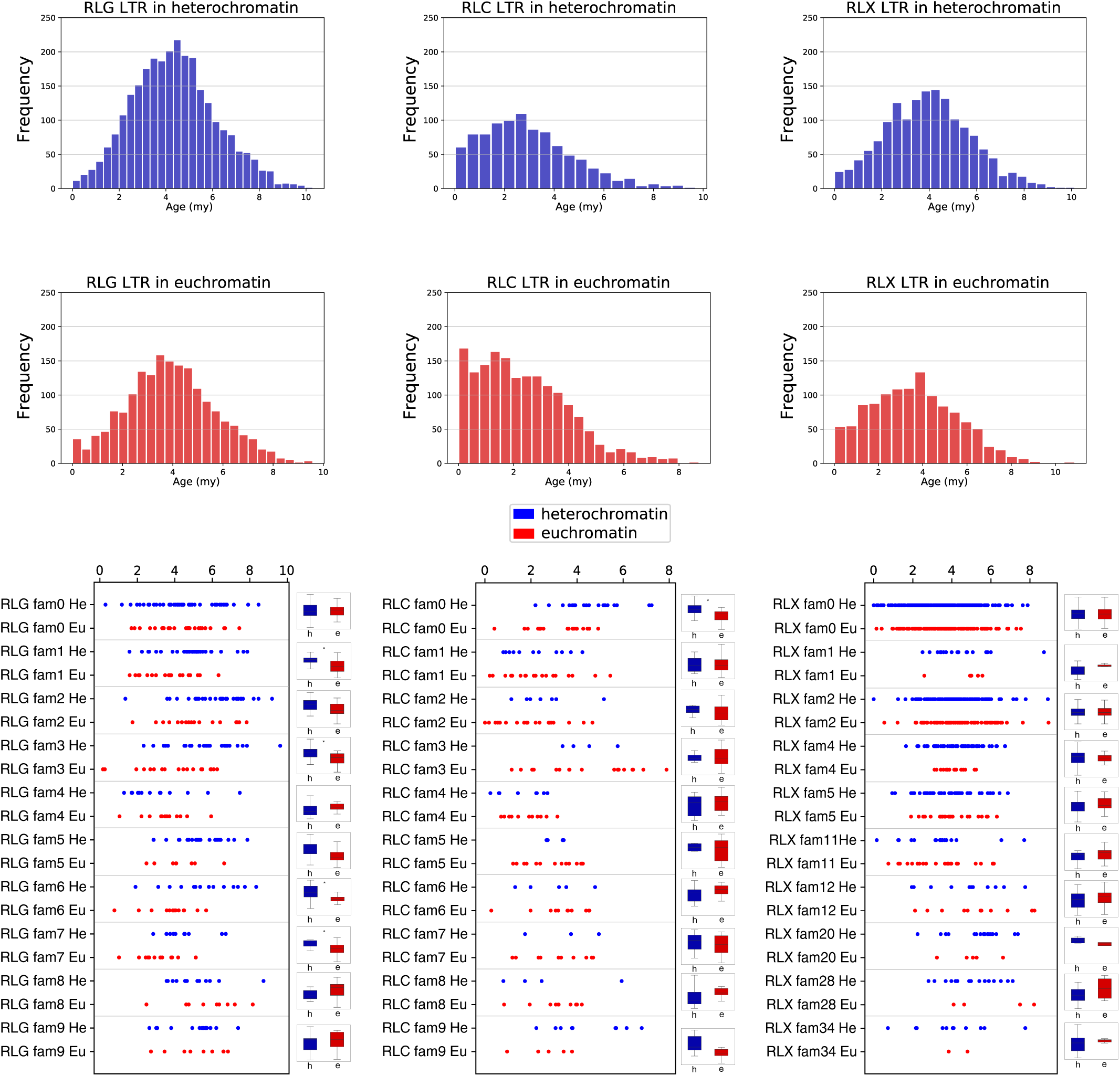
Number of LTR family by insertion age according to genome chromatin state. Upper panel: frequency histograms of Gypsy, Copia and Unclassified LTR families by their insertion age (in millions of years) separated by heterochromatin regions (blue) and euchromatic regions (red). Lower panel: Scatter plots of ten random independent families from Gypsy, Copia and Unclassified LTRs assessing the age distribution of individual elements by their heterochromatin (blue) and euchromatin (red) state. * shows significant differences between families (t-test p<0.05).

Furthermore, we plotted ten random independent families encompassing different member sizes from RLG, RLC and RLX to assess the age distribution of individual elements by region (Figure 5, bottom panel). Only four out of the ten evaluated Gypsy families showed significant differences (t-test p<0.05) in aging by region. In all the cases, heterochromatic elements were older than euchromatic elements, which reflect a slight age difference within elements of the same family owing to their insertion sites. This suggests that insertions into heterochromatic regions are more likely to persist for longer periods of time.

Overall, these variances may have a component based on differential mutation/substitution rate by region but more importantly it is strongly affected by superfamily traits. A recent study by Quadrana et al. (2019) described the mechanisms for which several TEs of the Copia superfamily preferentially integrate within genes by association with H2A.Z-containing nucleosomes. Moreover, they suggest that the role of H2A.Z in the integration of Copia retrotransoposons has been evolutionary conserved since the last common ancestor of plants and fungi (Quadrana et al., 2019).

On the other hand, Gypsy superfamily harbors a chromodomain that interacts with repressive histone marks such as H3K9m2 which targets to heterochromoatin (Sultana et al., 2017).

## Conclusion

TEs have been historically neglected in genome assembly projects, partially due to their repetitive nature but also because their heterogeneity in sequence, size, number of copies, distribution and mutation rates both inter and intra species, make them very difficult to detect accurately (Bourque et al., 2018).

However, in the past years, TEs discovery and annotation for the main crops have emerged with diverse results of coverage, complexity and accuracy. For instance, by using the CLARITE software on the IWGSC RefSeq v1.0 genome assembly, Alaux et al. (2018) found over 5 million elements from all types of TEs in wheat.

The Rice TE Database collects repeat sequences and TEs of several species of *Oryza* (rice) genus. All sequences have been characterized adopting Wicker’s classification code and extending it by encoding new TE superfamilies and non-TE repeats. Particular emphasis was given to the proper classification of sequences and to the removal of nested insertions (Copetti et al., 2015).

As we described above, potato has some TE sequences annotated on the RepBase, essentially, a list based on RepeatMasker provided by the Potato Genome Sequencing Consortium and the work from Mehra et al. (2015), which is based on RepeatModeler. However, this worldwide important crop still lacked a comprehensive, multiple-based discovery approach focused on transposon elements annotation.

This work arises as a need of a reliable potato TE atlas for ongoing projects that involve epigenetic and transcriptional regulation for different *Solanum tuberosum* in contrasting environments.

Plants are known for their phenotypic plasticity and good adaptation to environmental changes due to their sessile condition. There is an increasing evidence of the impact that TEs have on the transcriptome on response to stress. For example, TEs may have direct effects on genes regulation, by providing them new coding or regulatory sequences, changes on the epigenetic status of the chromatin close to genes, and more subtle effects by imposing diverse evolutionary constraints to different chromosomal regions (Makarevitch et al., 2015; Vicient and Casacuberta, 2017; Cambiagno et al., 2018). Finding common patterns or specific alterations on these features (TEs) and linking them with sRNAs profiles and DNA methylation patterns could help researchers to elucidate the underlying mechanisms of transcriptional changes during different stress conditions.

As mentioned above, the current data regarding TEs on potato genome is either scare or imprecise for transcriptional analysis. Given the importance of transposon elements, we provide here a comprehensive potato TE landscape, based on a wide variety of identification tools and approaches, clustering methods, copies detection, filtering rules and clear outputs, that the scientific community will likely use for metadata analysis.

## Acknowledgements

The authors would like to thank Dr. Soledad Lucero and Dr. Julia Sabio y Garcia for the assistance with English-language editing, and Humberto Julio Debat for his critical comments on the manuscript. This work was supported by the Instituto Nacional de Tecnología Agropecuaria (INTA) and by ANPCyT PICT 2015-1532, and PICT 2016-0429. The funders had no role in this study design, data collection and analysis, decision to publish or preparation of the manuscript.

## Author contributions

**DZ:** Conceptualization, Data curation, Formal analysis, Investigation, Methodology, Visualization, Writing - original draft, Writing - review & editing. **JMC:** Formal analysis, Data curation, Investigation, Methodology, Software, Writing – review & editing. **MG:** Investigation, Writing – review & editing. **ML:** Investigation, Writing – review & editing. **LSV:** Formal analysis, Investigation, Writing – review & editing. **RWM:** Funding acquisition, Investigation, Writing – review & editing. **SA:** Conceptualization, Funding acquisition, Investigation, Project administration, Resources, Supervision, Writing - review & editing.

## Electronic Supplementary Material

**Online Resource 1**. Fasta files containing family sequences for each type transposon elements.

**Online Resource 2**. gff3 file containing all TEs copy coordinates including Chr00.

**Online Resource 3**. Table resuming metrics of TEs to the nearest genes.

